# Conditions for the validity of SNP-based heritability estimation

**DOI:** 10.1101/003160

**Authors:** James J. Lee, Carson C. Chow

## Abstract

The heritability of a trait (*h*^2^) is the proportion of its population variance caused by genetic differences, and estimates of this parameter are important for interpreting the results of genome-wide association studies (GWAS). In recent years, researchers have adopted a novel method for estimating a lower bound on heritability directly from GWAS data that uses realized genetic similarities between nominally unrelated individuals. The quantity estimated by this method is purported to be the contribution to heritability that could in principle be recovered from association studies employing the given panel of SNPs 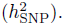 Thus far the validity of this approach has mostly been tested empirically. Here, we provide a mathematical explication and show that the method should remain a robust means of obtaining 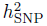 under circumstances wider than those under which it has so far been derived.

## Introduction

A central question in the study of quantitative phenotypic variation is the extent to which such variation is caused by genetic differences. The precise proportion of the phenotypic variance ascribable to genetic differences is formally known as the heritability. Many definitions of heritability have been proposed (Bell, 1977), but in this work we employ the *narrow-sense heritability* commonly denoted by *h*^2^ (Visscher et al, 2008). The concept of heritability was introduced by Fisher (1918) and Wright (1921) in their papers laying the foundations of quantitative genetics, although they did not use the word “heritability” in these early writings.

Its success notwithstanding, the imminent demise of quantitative genetics as a field of research has been repeatedly predicted ever since its first textbook appeared (Falconer, 1960; Hill and Mackay, 2004). Perhaps the prominent role in quantitative-genetic theory of heritability—a macroscopic parameter of a genetic system—has led some to suppose that advancing microscopic knowledge of the genetics underlying a given trait will superannuate the high-level approach. This anticipated obsolescence has not occurred, and indeed the recent explosion of findings from genome-wide association studies (GWAS) has only intensified the spotlight on the concept of heritability. For example, the loci found to be associated with a given trait at a strict threshold of statistical significance typically account for only a small proportion of the trait's heritability (as estimated from traditional studies of the correlations between close relatives), and this discrepancy has led to much discussion of “missing heritability” (Manolio et al, 2009; Eyre-Walker, 2010; Dickson et al, 2010; Wray et al, 2011; Zuk et al, 2012; Gibson, 2012; Hemani et al, 2013).

The estimation of heritability from the correlations between relatives has been substantially augmented by a novel technique that makes use of dense GWAS data from nominally unrelated individuals (Yang et al, 2010; Visscher et al, 2010; Lee et al, 2011). This technique is perhaps the most important innovation in quantitative genetics to have been introduced in the last dozen years, and it has provided what some may regard as decisive evidence for the view that undiscovered common variants account for a substantial portion of missing heritability. We will follow Benjamin et al (2012) and refer to this method as *genomic-relatedness-matrix restricted maximum likelihood* (GREML).

The descriptions of GREML given in the literature suggest that the mathematical basis of this method is not fully understood. For instance, formal justifications of the method that have been offered so far seem unable to account for biased estimates occasionally observed in simulation studies. Here we attempt to further the mathematical understanding of GREML, thereby providing insight into cases where the method has not worked well. By revealing the sense in which these cases are extreme, however, our account conversely shows that GREML estimates are in fact quite robust. We treat the heritability of a single trait, but our account can be generalized to the genetic correlation between two traits.

## Subjects and methods

We emphasize here that some of our mathematical arguments employ restrictive assumptions about sample size, the number of genotyped markers, and the values of the variance components. However, the fact that certain assumptions are sufficient to prove a result does not imply that the assumptions are necessary, and later we provide strong evidence for the generality of our findings.

We illustrate some of our mathematical arguments with numerical simulations, using two GWAS datasets to supply the genetic data. One dataset was used in a GWAS of European Americans reported previously (Chabris et al, 2013). The quality-control filters left 401 individuals and 661,108 markers (although only subsets of markers on chromosome 1 were used). We employed this small-sample dataset when it was necessary to relieve computational burden.

The second dataset was taken from the GENEVA Genes and Environment Initiatives in Type 2 Diabetes (Nurses’ Health Study/Health Professionals Follow-Up Study). We used PLINK to eliminate individuals of reported non-European descent, markers missing more than 5 percent of their calls, markers showing significant deviation from Hardy-Weinberg equilibrium (HWE) (*p* < 1 × 10^−6^), markers with minor allele frequency (MAF) *<* 0.01, individuals missing more than 5 percent of their genotypes, and one individual from any pair with a relatedness (Eq. 5) exceeding 0.025 in absolute value; Zaitlen et al (2013) provide some discussion of the appropriate relatedness cutoff. These filters left 4,975 individuals and 697,709 markers.

We used the software tool LDAK to calculate the extent to which each SNP is tagged by its neighbors (Speed et al, 2012). In particular, we computed the matrix

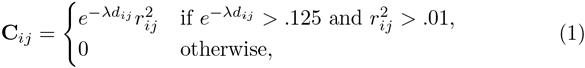

where *d_ij_* is the distance between SNPs *i* and *j* in base pairs (equaling ∞ if the SNPs are on different chromosomes), 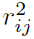 is the standard measure of linkage disequilibrium (LD) between *i* and *j*, and *λ* is chosen so that exp(−*λd_ij_*) = .125 when *i* and *j* are 3 Mbp apart. SNP *i*'s level of tagging by its neighbors is then the sum of the elements in the *i*th row of **C**. Large values of **C** correspond to strong tagging (redundancy), whereas small values correspond to a lack of LD with neighbors.

Table 1 lists the markers used in the simulations testing the case of a single marker with a nonzero partial regression coefficient. The mean of 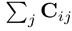 over all *i* was approximately 11.45 and the standard deviation approximately 8.35, and we chose three SNPs with values close to the mean as the “moderately tagged” SNPs. Similarly, we choose three SNPs close to the 3rd percentile (2.05) as the “very weakly tagged” SNPs, three SNPs close to the 20th percentile (5.02) as the “weakly tagged” SNPs, three SNPs close to the 80th percentile (16.58) as the “strongly tagged” SNPs, and three SNPs close to the 97th percentile (30.36) as the “very strongly tagged” SNPs. Within each group of three markers, one was chosen to have an MAF of ∼ 0.01, another to have an MAF of ∼ 0.25, and the last to have an MAF of ∼ 0.50. More specifically, for all markers within a given percentile of the 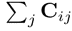 distribution plus/minus 0.05, one random selection was made from the markers with MAF in the interval (0.01, 0.02), another from markers in the interval (0.245, 0.255), and yet another from the interval (0.49, 0.50) to create a set of three markers varying in MAF but matched with respect to LD. The extent of tagging by neighbors is moderately correlated with MAF, and there were no candidates for very strongly tagged markers meeting the initial requirement for low MAF. The right endpoint of the low-MAF interval was therefore extended by increments of .01 until the set of candidates was nonempty.

We used GCTA to simulate phenotypes and estimate heritabilities on the basis of GREML. Each simulation scenario was tested with 200 replicates.

## Results

Consider a sample of *n* unrelated (very distantly related) individuals and *p* biallelic markers. Let 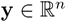 be the vector of standardized phenotypes, 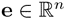 the vector of residuals (the sum of non-additive genetic deviations, environmental deviations, and errors of measurement), 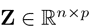 the matrix of standardized genotypes, and 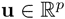 the vector of partial regression coefficients in the regression of the phenotype on standardized genotypes. If *X_ik_* is the count of minor alleles (0, 1, or 2) carried by individual *i* at marker *k*, then HWE implies that the standardized count is 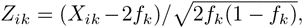 where *f_k_* is the MAF at marker *k*. Note that the elements of **u** are not necessarily proportional to the average effects of gene substitution (Fisher, 1941; Lee and Chow, 2013), since a non-causal marker may have a nonzero coefficient because it is in LD with a causal locus that has not been genotyped. The phenotype need not be standardized, but it makes our presentation simpler.

The residuals will be uncorrelated with the “chip-based” breeding (additive genetic) values **Zu** if gene-environment correlation is absent or properly controlled (Yang et al, 2011b; Browning and Browning, 2011; Goddard et al, 2011; Janss et al, 2012; Speed et al, 2012) and if **Zu** and the vector of genotypic means are orthogonal. What the latter condition means is that the chip-based breeding values in **Zu** must be uncorrelated with both the discrepancy between true and chip-based breeding values and the non-additive residuals attributable to dominance and epistasis. This condition is difficult to assess, but we assume henceforth that it is met. Then from the basic equation

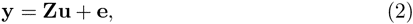

we see that the total phenotypic variance can be written as

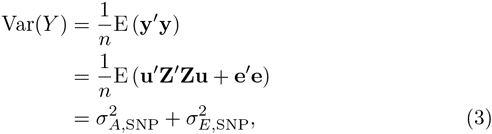

where the expectation is over random residuals. The “SNP-based heritability” is thus 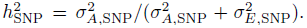 If we assume that linkage equilibrium (LE) holds approximately, then **Z′Z** ≈ *n***I***_p_* and the additive genetic variance is approximately **u′u**.

We emphasize that 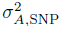 is the variance that would be removed from the total phenotypic variance by multiple regression on all markers that happen to be assayed by the genotyping chip, as sample size goes to infinity. Because not all causal variants may be genotyped or represented by LD proxy, the chip-based additive genetic variance denoted here by 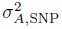 is smaller than the true additive genetic variance 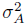 contributed by all causal loci. Similarly, 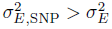 and 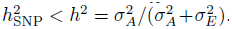 Leaving aside these subtleties of definition, we can see that Eq. 3 holds because (1*/n*)E (**u′Z′Zu**) is the variance of chip-based breeding values and hence equal to 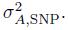

GREML estimates the parameters 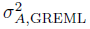 and 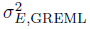 in the model

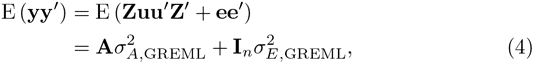

where, in the notation of Yang et al (2010), **A** = (1*/p*)**ZZ′** is the matrix of realized relatedness coefficients. It is helpful to write out the typical element of **A**,

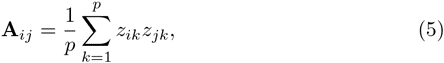

which is very analogous to the traditional coefficients of relatedness appearing in the classical formulas for the correlations between close relatives (Crow and Kimura, 1970; Lynch and Walsh, 1998). Preserving this analogy turns out to be important because different possible standardizations of the *X_ik_* will lead to different estimates of the variance components. For example, if a matrix differing from **A** by a constant factor is used in the place of **A**, then the estimate of 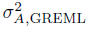 will be multiplied by that constant (Speed et al, 2012). The average of the off-diagonal realized relatedness coefficients over all pairs is zero (Powell et al, 2010), and the diagonal elements of **A** converge to unity as *p* becomes large.

At this point the equality of the first and second lines in Eq. 4 should be regarded not as a derivable fact but rather as *a priori* definitions of the parameters 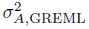 and 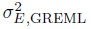. Note that 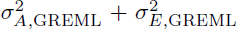 = 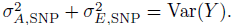

The emergence of the matrix **A** from the action of the expectation operator in Eq. 4 implies that **A** is a constant (up to permutations of the sample) characterizing the population from which the *n* individuals have been drawn. The expectation must thus be interpreted as taken over random samples of size *n* sharing the same precise histogram of relatedness coefficients **A***_ij_* but differing in the specific entries of **e**. In this way **A** is somewhat analogous to the sum of the squared differences between the independent variable and its mean (an ancillary statistic) in Fisher's (1973) discussion of univariate linear regression. Since fixing **Z** suffices to fix **A**, we henceforth assume that **Z** is fixed. This interpretation does not seem problematic; across distinct samples of large size *n* from the same population, genotyped with the same *p*-variant chip, histograms of relatedness coefficients with a reasonable shared bin width should exhibit little variability.

Let us compare Eqs. 3 and 4. The expectation of **ee′** alone is 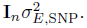 Therefore, 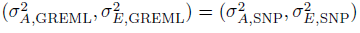 implies that **Zuu′Z′** = 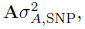 which in turn implies that **Zuu′Z′** is proportional to **u′uZZ′**. Such proportionality does not hold, however, as a matter of mathematical necessity. Therefore the question is posed: under what circumstances is

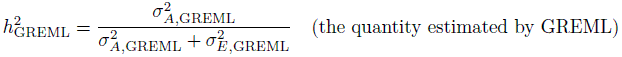

approximately equal to

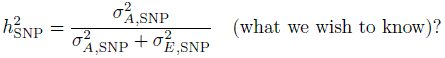

It is important to note that no assumption-free adjustment of 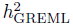 can yield a reliable estimate of the true heritability *h*^2^ if the genotyping chip assays a limited panel of markers. For example, it may be that rare causal variants have large phenotypic effects, and such variants may be absent from the genotyping chip and poorly represented by LD proxy. Therefore, if it turns out that 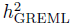 bears no exact relationship to *h*^2^, this should not necessarily be construed as a fault of GREML. The most that can be reasonably demanded from the method is that 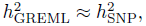 and this is the issue that we address here.

Yang et al (2010) assume that each element of **u** can be regarded as an independent draw from a normal distribution with mean zero and variance 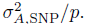 The desired equality between 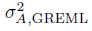 and 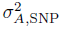 then follows if we further suppose that the treatment of **u** as a vector of independent random variables justifies the replacement of **uu′** with 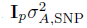 under the action of the expectation operator. There are two aspects of this assumption, however, that seem rather nonbiological. The first is that the number of markers with nonzero regression coefficients (“nonzeros”) is typically believed to be much smaller than the total number of genotyped markers (Park et al, 2011; Stahl et al, 2012), which is inconsistent with a normal distribution. Secondly and more importantly, the partial regression coefficients in **u** represent the average effects of gene substitution (or LD proxies for such effects) and thus cannot be said to vary randomly across individuals. Hence, while the spectrum of the coefficients could be described by a normal distribution or some other distribution, the exterior product **uu′** cannot be averaged over this distribution characterizing *markers* when given as an input to an expectation operator over random residuals disturbing the phenotypes of *individuals*. This implies that **uu′** is not proportional to the identity matrix.

Note the contrast between the meaning of **u** in the GREML literature and in the treatment of linear mixed models by Lynch and Walsh (1998). For instance, in the example of Lynch and Walsh's Chapter 26, the elements of **u** are the breeding values of the pedigree founders and thus can properly be said to vary randomly across different realizations of the pedigree structure.

For the reasons just given, we refrain from the Yang et al (2010) assumption and treat **u** as an arbitrary fixed constant rather than a random variable. Fixing both **Z** and **u** implies that the population targeted for inference consists of random samples sharing the same *sample* value of 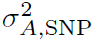 (Eq. 3). This limitation does not seem unduly restrictive since, for values of *n* often used in GREML applications (∼10,000), different samples from the same population (e.g., Northwest Europeans) will scarcely differ in their realized values of 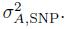

We can now write the typical off-diagonal element of the matrix **Zuu′Z′** in component form as

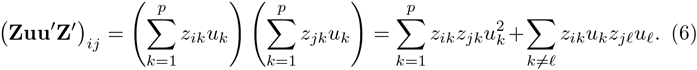

Now suppose that the constants 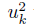 were all equal to 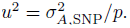 Then the first sum in the expression above would become 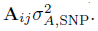 However, since it is surely false that each marker has the same squared coefficient in the regression on standardized genotypes, we decompose each 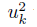 into the sum 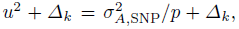 where 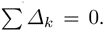 If there are many genotyped markers, then *u*^2^ is a rather small quantity. If the nonzeros are a small fraction of the total, then most of the *Δ_k_* are equal to *−u*^2^. If the *k*th marker has a nonzero coefficient, however, then its *Δ_k_* has a relatively large positive value.

Then we have, for the typical off-diagonal element of **Zuu′Z′**,

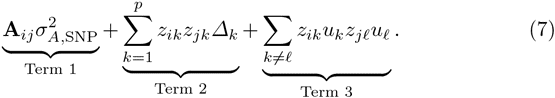

We have already stated that the sum of Term 1 over all pairs of individuals is zero (i.e., the sum of **A***_ij_* over *i* ≠ *j* is zero). Since the *Δ_k_* are constants, the sum of Term 2 over all pairs is also zero. The sum of Term 3 over all pairs is zero as well because knowledge of randomly chosen individual *i*'s genotype at marker *k* cannot provide any information about randomly chosen individual *j*'s genotype at marker ℓ, even if *k* and ℓ are in perfect LD; only individual *j*'s own genotype at *k* can provide this information.

It might seem that Terms 2 and 3 must be extremely small compared to Term 1 for each pair of individuals *i* and *j* in order for 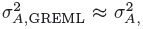 to hold. In genetic data, however, this condition will rarely be fulfilled. To illustrate this fact, we simulated 1,000 individuals with genomes consisting of 100 markers in LE and MAFs drawn uniformly from (0.05, 0.50). The elements of **u** were drawn from a normal distribution and constrained to produce 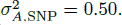 The results of calculating all three terms in Eq. 7 for each pair show that Term 2 is often comparable in magnitude to Term 1 and that Term 3 is frequently larger (Fig. 1). Increasing the number of markers from 100 to 1,000 only reinforced this conclusion. It is also of some interest to calculate the three terms in Eq. 7 using real data so as to examine the impact of LD. We therefore used the Chabris et al (2013) data to run simulations using distinct sets of 100 and 200 contiguous SNPs on chromosome 1. Each real SNP typically showed moderate to strong LD with neighbors, and there were pairs of SNPs with values of *r*^2^ exceeding 0.90. As can be seen in Fig. 2, the use of real data did not cause Terms 2 and 3 to vanish. Note that the average of each term was still close to zero.

**Fig. 1.**
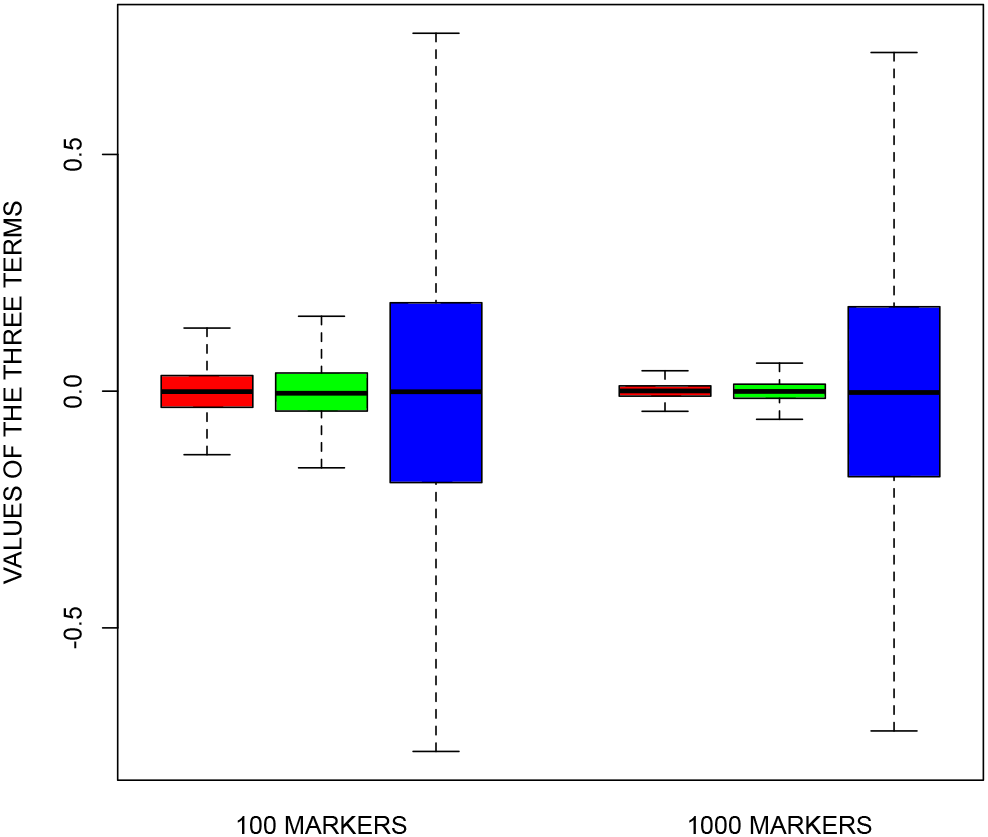
Boxplots of the three terms in Eq. 7 for pairs of simulated individuals and **u**. Red corresponds to Term 1, green to Term 2, and blue to Term 3. The plots on the left display the results for 100 markers in LE, whereas those on the right display the results for 1,000 markers in LE. Values lying beyond 1.5 times the interquartile range past either the 25th or 75th percentile are omitted. Note that the average of each term is close to zero.

**Fig. 2.**
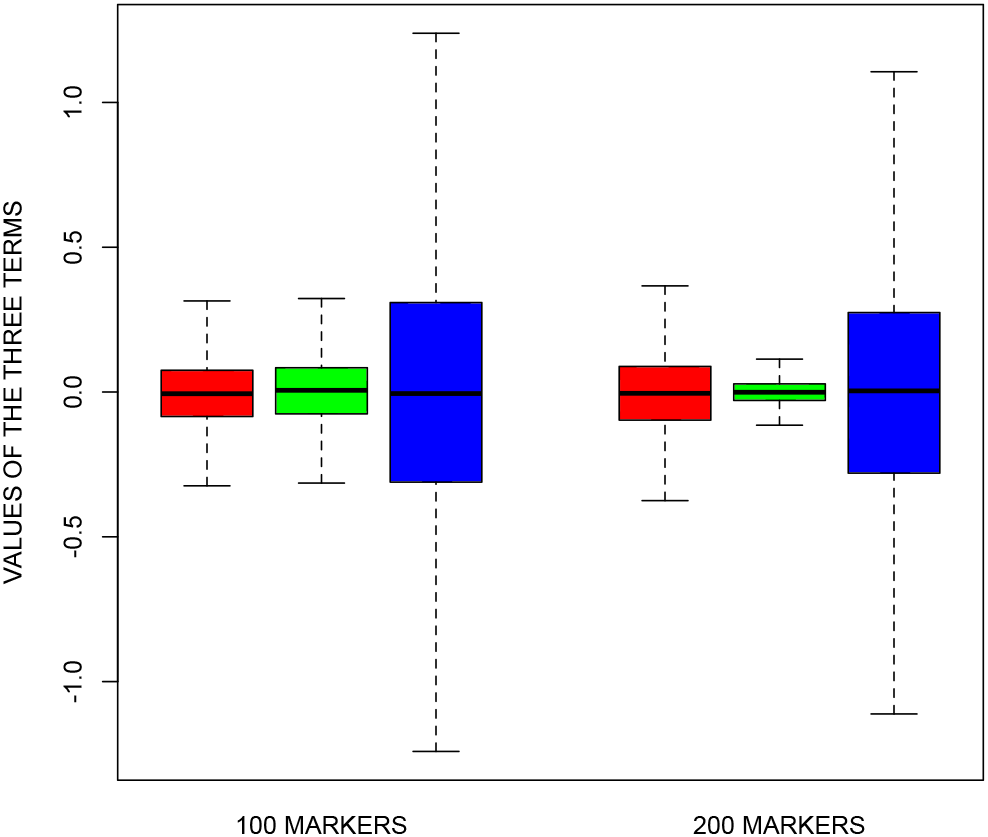
Boxplots of the three terms in Eq. 7 for pairs of real genotyped individuals (but simulated **u**). Red corresponds to Term 1, green to Term 2, and blue to Term 3. The plots on the left display the results for 100 contiguous genotyped markers on chromosome 1, whereas those on the right display the results for 200 such markers elsewhere on the same chromosome. Values lying beyond 1.5 times the interquartile range past either the 25th or 75th percentile are omitted.

Despite the failure of Terms 2 and 3 to vanish, GREML is often unbiased as a means of estimating 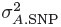 when applied to real genetic data (Speed et al, 2012, 2013; Zhou et al, 2013; Browning and Browning, 2013; Lee et al, 2013), which implies that the vanishing of the additional terms is not a necessary condition. Here we seek a more general characterization of those cases where GREML is accurate.

The variance components are estimated with GREML using REML (Lynch and Walsh, 1998; Yang et al, 2010, 2011a; Vattikuti et al, 2012). We now derive certain conditions that the maximum-likelihood (ML) estimates must satisfy. The GREML model in Eq. 4 is equivalent to treating **y** as drawn from a multivariate normal distribution with mean zero and covariance matrix **V** = 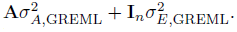 In the absence of fixed effects, the log-likelihood is thus

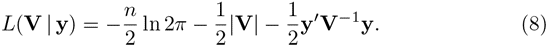

The ML estimates of the variance components are obtained by taking the partial derivatives of Eq. 8 with respect to 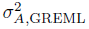 and 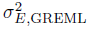 and setting the resulting equations to zero. Now recall that if **M** is a square matrix whose elements are multiples of a scalar *x*, then

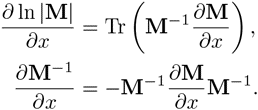

Using these facts, we differentiate Eq. 8 and obtain

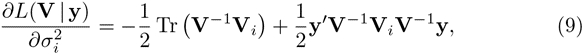

where

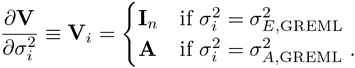

The ML conditions are thus

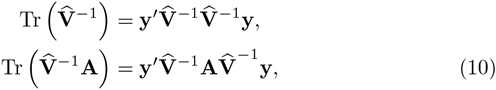

where 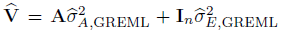 is the ML estimate of **V**. Since ML estimates are consistent given mild regularity conditions, we set the estimate 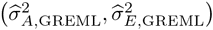 equal to 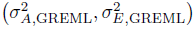 for convenience.

To express the GREML variance components in terms of the observables **y** and **A** (which in turn depend on the parameters of primary interest 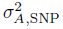 and 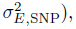 we need an approximation for **V**^−1^. A standard linear algebra approximation of the matrix inverse is

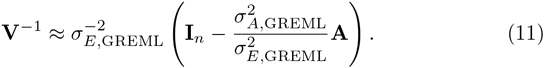

Multiplying Eq. 11 by **V**, we obtain

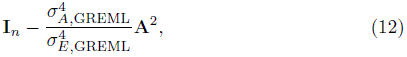

and thus the vanishing of the second term will render Eq. 11 a good approximation of **V**^−1^. Given *p* markers in LE, the average of the squared off-diagonal elements of **A** is close to 1/*p* (which we confirmed in the simulations generating Fig. 1). The typical off-diagonal element of **A**^2^, 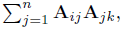 is thus likely to be smaller than *n/p*. It follows that 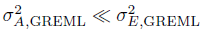 and *n* ≪ *p* are sufficient conditions for Eq. 11 to serve as an approximation of **V**^−1^. When Eq. 11 is substituted into Eq. 10, we obtain

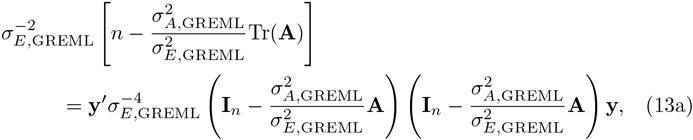

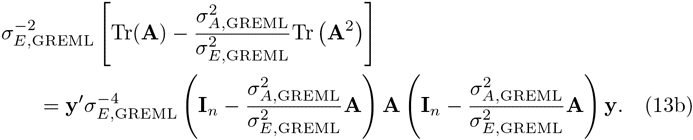

In principle, we need to solve this system of two equations with two unknowns. However, for large *n* and a standardized phenotype we can immediately impose the equality 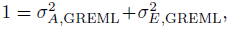 and noting that Tr(**A**) ≈ *n*, we obtain the single equation

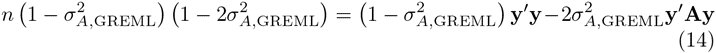

to first order, which in turn implies

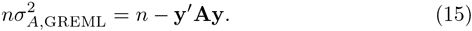

Therefore, if 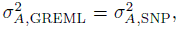 then 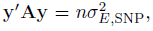 which corresponds to a necessary condition for GREML to provide a consistent estimator of 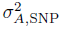 in the small-(*n/p*), small-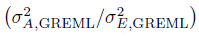 regime.

We now examine conditions under which 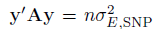 holds. Let us use **S***_ij_* to denote the sum of Terms 2 and 3 in Eq. 7. Then the expectation over random residuals of the quadratic form **y′Ay** can be written as

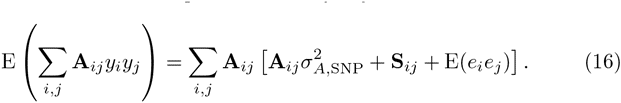

The last term is indeed 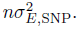 The first term, 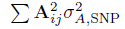, has a diagonal contribution converging to 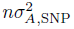 and an off-diagonal contribution converging to 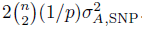 Therefore the first term is approximately (*n* + *n*^2^/*p*)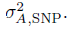 The ratio of the first and last terms is approximately (1 + *n*/*p*)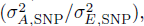 and thus the sufficient conditions for Eq. 11 to give **V**^−1^ also ensure that the contribution of the last term to Eq. 16 dominates that of the first.

The second term in Eq. 16, 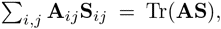 must also be close to zero for 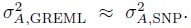 The diagonal contribution to this sum, 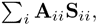 converges to 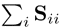 as *p* becomes large. Since **S***_ii_* is the deviation of individual *i*'s squared breeding value from 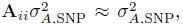 the sum of these deviations over all individuals becomes zero. The off-diagonal contribution to Tr(**AS**) can be interpreted as (proportional to) a covariance between relatedness **A***_ij_* and the sum of Terms 2 and 3, and this covariance must also be zero.

The sign of Tr(**AS**) when it deviates from zero cannot in general be used to predict the sign of 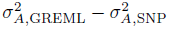 from Eq. 15, which is derived from an uncontrolled expansion with an unknown range of validity. The following argument for the direction of the bias induced by Tr(**AS**) ≠ 0 appears to be valid for typical values of *n*, *p*, and 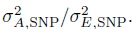 If there is a positive covariance between **A***_ij_* and **S***_ij_*—meaning that pairs with above-average (below-average) relatedness also tend to have above-average (below-average) values of Terms 2 or 3—the phenotypic products *y_i_y_j_* are systematically too far from zero and thus lead GREML to infer an excessive SNP-based heritability 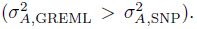 Conversely, if there is a negative covariance between Term 1 and the sum of Terms 2 and 3, the shrinking of phenotypic products toward zero leads GREML to underestimate SNP-based heritability 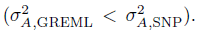 There is a close analogy here to the requirement of a zero correlation between the causal variable and the residual disturbance for least-squares regression to provide an unbiased estimate of a linear causal effect.

Note again that we have made no assumption with respect to whether the partial regression coefficients in **u** follow a normal distribution or indeed any probability distribution. In fact, as we will shortly demonstrate, GREML can serve as an accurate means of estimating 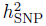 in the case of a single nonzero, and obviously a probability distribution prescribing one nonzero and *p* − 1 zeros is not normal. Therefore this feature of the genetic architecture *per se* should not affect the accuracy of GREML as a method for estimating 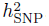.

We now show that our account provides quantitative explanations of recent simulation results. Both Speed et al (2012) and Zhou et al (2013) remarked upon the fact that GREML remains approximately unbiased even as the number of nonzeros becomes very small. This is perhaps surprising because the majority of the *Δ_k_* in this case are equal to 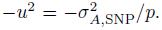 But suppose that there are *s* nonzeros, where *s* is an arbitrary positive integer smaller than or equal to *p*. As long as the markers are in LE, then the contribution to Tr(**AS**) from the products of relatedness and Term 2 is zero since

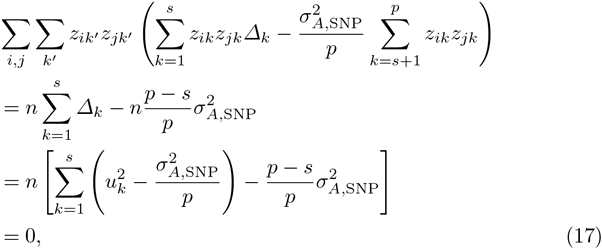

where we used the property that LE and large *n* imply 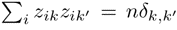 (δ_k,k′_ is the Kronecker delta). Note that each of the *s* nonzeros may have an arbitrary MAF and *u_k_*. Furthermore, the typical term in the expansion of the covariance between relatedness and Term 3 is

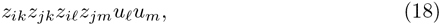

and LE also ensures that the average of this product vanishes. Whenever *ℓ* or *m* indexes a marker outside of the nonzeros, the product vanishes regardless, and a single nonzero thus trivially guarantees a zero covariance. Therefore, since LE guarantees that Tr(**AS**) = 0 in the small-(*n/p*), small-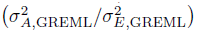 regime, GREML will accurately estimate the heritability captured by a set of independent markers even as the number of nonzeros decreases down to one.

It has been reported that the MAF spectrum of the nonzeros can affect the accuracy of GREML (Speed et al, 2012, 2013; Lee et al, 2013). Because the calculations related to Eqs. 17 and 18 rely on LE rather than any assumption regarding the MAF spectrum, this sensitivity must arise from LD and the tendency of higher-MAF variants to be better tagged by neighboring markers. One way in which LD affects GREML can be easily explained upon making the convenient assumption that the panel of markers is partitioned into two subsets, one of which is characterized by complete LE and the other by complete LD. Then Term 2 can be rewritten as

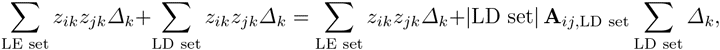

where |·| denotes set cardinality and **A**_*ij*,LD set_ is the relatedness over just the markers in the LD set. The factor can be pulled out from the second sum because perfect LD implies that *z_ik_z_jk_* equals the constant |LD set| **A**_*ij*,LD set_ for each *k* in the set. Notice that **A**_*ij*,LD set_ makes a disproportionate contribution to **A***_ij_*. For example, if there are 10,000 markers and |LD set| = 5,000, then **A***_ij_* is a weighted sum of 5,001 contributions where the weight of the LD set is 5,000 times as large as any other. If the nonzeros (positive *Δ_k_*) are all in the LD set, then a positive correlation may be induced between Terms 1 and 2. Conversely, if the nonzeros are all in the LE set, then a negative correlation may be induced because *Δ_k_* = − *u*^2^ for each *k* in the LD set. A small number of nonzeros can therefore lead to upward (downward) bias because by chance the nonzeros may be strongly (poorly) tagged by neighboring SNPs. On the other hand, suppose that there are an equal number of nonzeros in the LE and LD sets. So long as there is no tendency for the *Δ_k_* of the nonzeros to be larger in one of the sets, the magnification of the nonzeros in the LD set should be balanced by the diminution of those in the LE set, leading to an overall estimate of 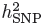 that may be nearly unbiased.

To illustrate the impact of LD on the covariance between relatedness and Term 2, we performed simulations where each of 1,000 contiguous SNPs on chromosome 1 in the Chabris et al (2013) data was stipulated to be the single nonzero. Fig. 3 shows that there was a strong correlation, approaching unity, between the chosen nonzero's redundancy with neighboring SNPs (Eq. 1) and the resulting covariance between relatedness and Term 2. Because we used only a single nonzero in each simulation, Term 3 was trivially zero for all pairs and therefore did not need to be computed. The results displayed in Fig. 3 clearly bear out the fact that a clustering of nonzeros among the most (least) well-tagged markers leads to a positive (negative) covariance between relatedness and the additional terms in Eq. 7.

**Fig. 3.**
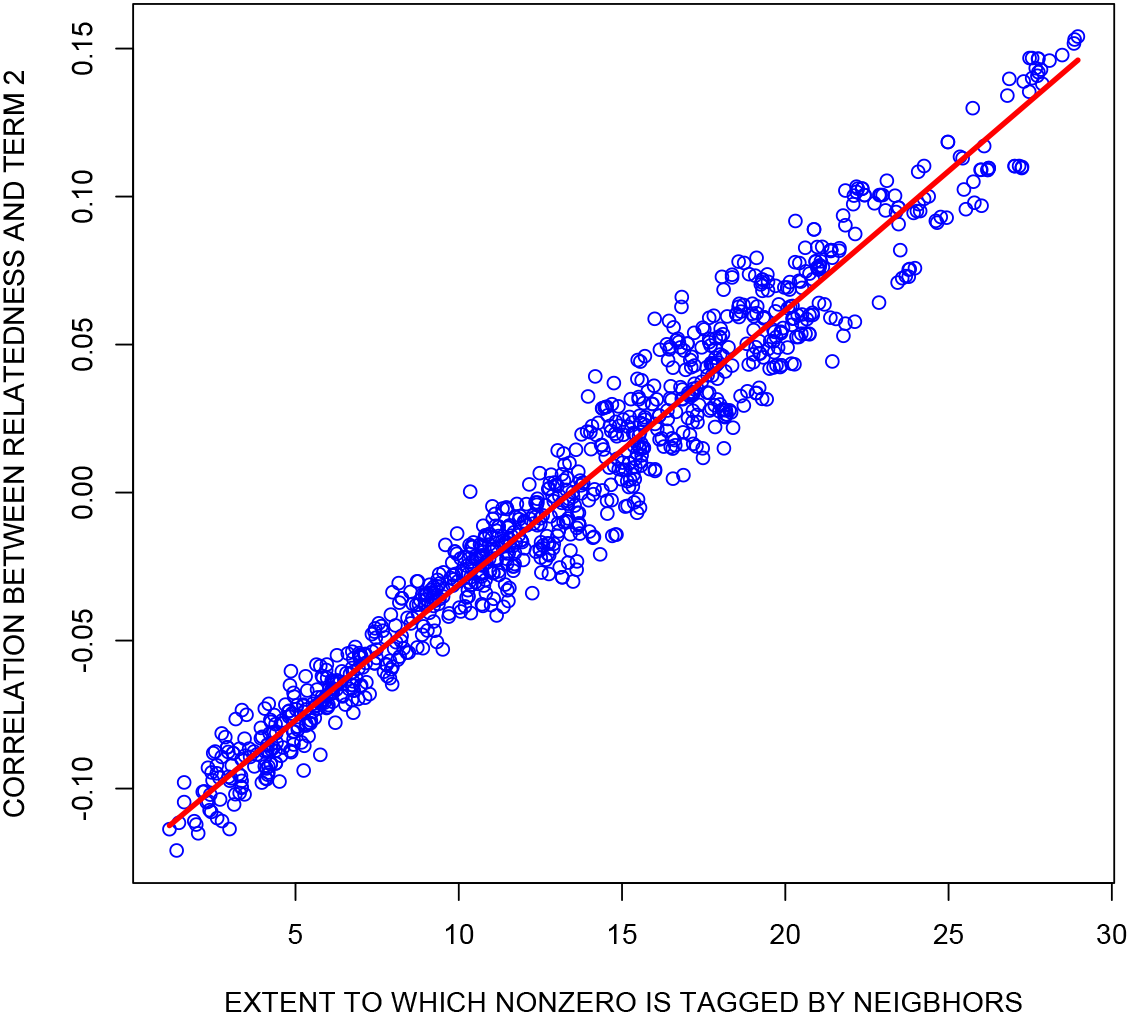
The strong relationship between the extent to which the simulated nonzero SNP is tagged by neighboring markers and the resulting correlation between relatedness (Eq. 5) and Term 2 (Eq. 7). Each point represents one of 1,000 contiguous genotyped SNPs on chromosome 1 in the dataset of Chabris et al (2013). The *x*-axis represents the sum of the elements in the row of **C** (Eq. 1) corresponding to the nonzero SNP, and the *y*-axis represents the correlation between relatedness and Term 2. The LOESS curve is displayed in red.

A shortcoming of our mathematical expressions is that they make no predictions when either *n/p* or 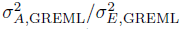 is relatively large. However, because the requirements of small *n/p* and 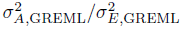 only arise from our need to approximate **V**^−1^ for the purpose of obtaining a closed-form solution of the ML equations, it is quite plausible that our condition for the unbiasedness of GREML as a method for estimating 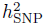 continues to be necessary outside of the small-(*n/p*), small-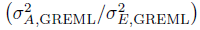 regime. In particular, the condition of a zero correlation between relatedness and the additional terms in Eq. 7 should continue to hold for the following intuitive reason. If we reduced the dataset such that the phenotypic products of pairs within a given bin of relatedness (plus/minus a small quantity) were averaged together, then the conditional average of the additional terms equaling zero given any relatedness would ensure that the average phenotypic product of pairs within a fixed bin of relatedness is equal to that relatedness times 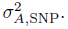

We found that the small values of *n*, *p*, and *s* used in our simulations based on the Chabris et al (2013) dataset prevented the diagonal contribution to Tr(**AS**) from closely approaching zero, thereby rendering this dataset unsuitable for simulations extrapolating our deductions. For this reason we turned to our second dataset, where *n* and *p* are more typical of GREML applications. In this series of simulations, each of the markers in Table 1 was specified in turn to be the single nonzero. The true 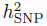 was set to 0.50, and we recorded the 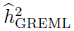 produced by each replicate.

**Table 1.** SNPs used in simulations of GREML performance in the case of one nonzero

| SNP $i$ | chromosome | MAF ($f_i$) | $\sum_j \mathbf{C}_{ij}$ |
| --- | --- | --- | --- |
| *very weakly tagged nonzeros* |  |  |  |
| rs4674229 | 2 | 0.0119 | 2.08 |
| rs4716447 | 7 | 0.2460 | 2.02 |
| rs4968679 | 17 | 0.4978 | 2.09 |
| *weakly tagged nonzeros* |  |  |  |
| rs11039838 | 11 | 0.0108 | 5.01 |
| rs4912830 | 5 | 0.2514 | 5.04 |
| rs2654534 | 15 | 0.4955 | 5.02 |
| *moderately tagged nonzeros* |  |  |  |
| rs7620645 | 3 | 0.0176 | 11.49 |
| rs12692474 | 2 | 0.2526 | 11.42 |
| rs4870308 | 6 | 0.4915 | 11.41 |
| *strongly tagged nonzeros* |  |  |  |
| rs16841231 | 2 | 0.0156 | 16.54 |
| rs6424728 | 1 | 0.2505 | 16.61 |
| rs328890 | 7 | 0.4926 | 16.58 |
| *very strongly tagged nonzeros* |  |  |  |
| rs10138824 | 14 | 0.0278 | 30.36 |
| rs2718306 | 7 | 0.2521 | 30.37 |
| rs8006587 | 14 | 0.4910 | 30.34 |

The results displayed in Fig. 4 bear out our predictions.

1. It is possible for 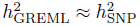 even if there is only one nonzero, as long as its level of LD with neighbors is typical of the SNP panel. In the case of a moderately tagged nonzero, no average 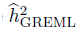 missed the true 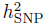 by more than 0.014.
2. Strong (weak) tagging leads to upward (downward) bias in 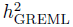 as an estimate of 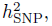 and this bias increases with the nonzero's deviation from the typical level of tagging. This dose-response relationship is consistent with the increasing magnitude of the correlation between relatedness and Term 2 in Eq. 7 as the nonzero deviates from the typical level of tagging (Fig. 3). There was one anomalous result: our randomly chosen marker satisfying our criteria for low MAF and poor tagging (rs11039838) showed an average 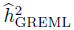 of 0.487, not far from the true 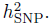 To determine whether this was an unusual deviation from the overall trend, we reran the simulation of this scenario, this time specifying all 125 markers satisfying our criteria for low MAF and poor tagging as nonzeros of equal coefficient magnitude (*u_k_*). The resulting average 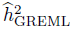 was 0.347 [95% CI = (0.340, 0.353)], further from the true 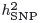 and closer to those observed at the other MAFs within our group of poorly tagged markers.
3. Once the level of tagging is controlled, the MAF of the nonzero has no discernible systematic impact on 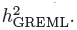 Even the anomalous result produced by our initial choice of a poorly tagged low-MAF marker (rs11039838) deviated in the opposite direction from the prediction of an account positing an association between low MAF of a nonzero *per se* and underestimation of 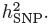
4. The average of the heritability estimates across all 15 markers (0.525) was close to the true 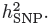

**Fig. 4.**
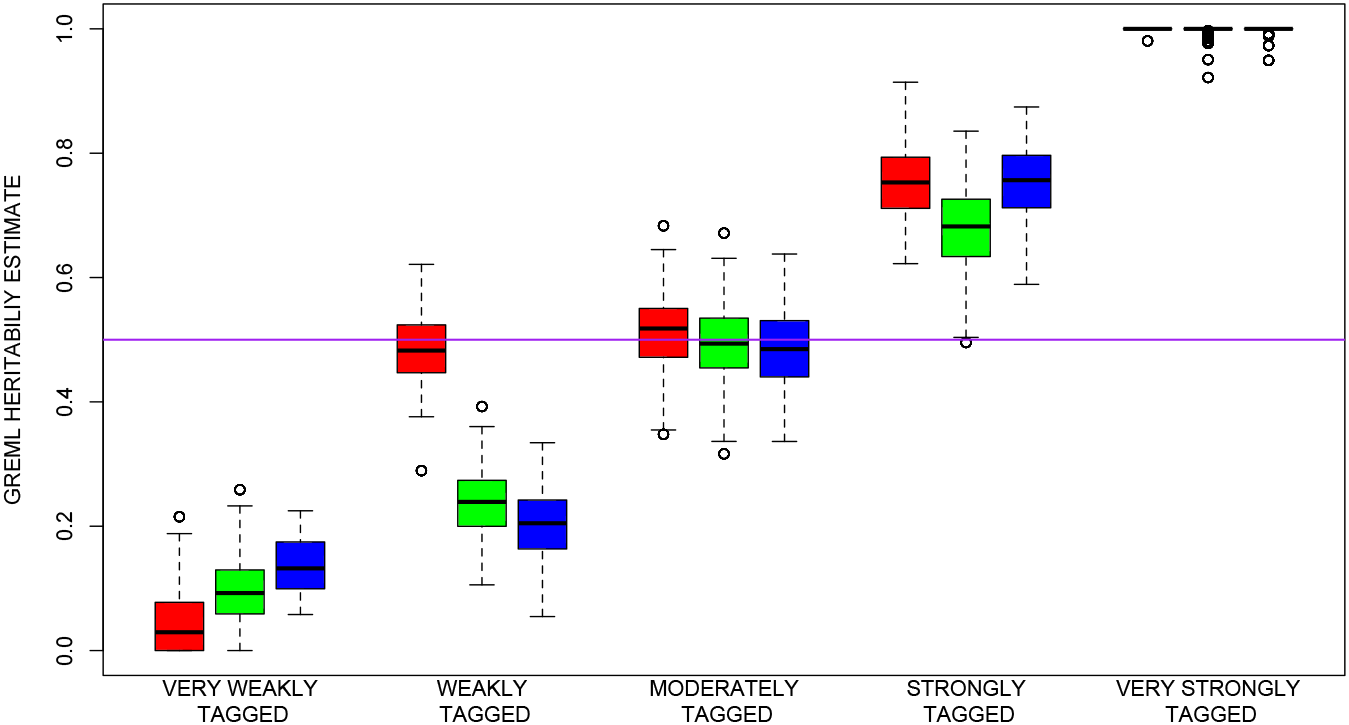
Simulations of GREML performance in the case of a single nonzero. Each group of three markers was characterized by very similar tagging levels within the group. Red corresponds to markers with MAF ∼0.01, green to MAF ∼0.25, and blue to MAF ∼0.50. Each scenario was tested with 200 replicates, which led to very precise estimates of the central tendencies. The purple horizontal line corresponds to the true 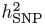 of 0.50.

To confirm that 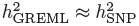 if the nonzeros are representative of the entire genotyping chip with respect to tagging, we ran another simulation specifying 10,000 randomly chosen markers as the nonzeros. The distribution of tagging (Eq. 1) in this subset was nearly identical to the distribution among all 697,709 genotyped markers. The mean of the 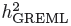 estimates was 0.499 [95% CI = (0.487, 0.511)]. It is worth pointing out that we drew the magnitudes of the nonzero coefficients from a normal distribution. If the distribution is such that a few coefficients are much larger than others, than it is possible that chance unrepresentative tagging of the dominating nonzeros will lead to some bias. However, this potential problem does not seem too threatening in practice, because loci of large effect are relatively easy to detect and can be removed from the analysis.

Interestingly, although we have not mathematically analyzed the standard errors produced by GREML software, we found in our simulations that GCTA's standard errors are reasonably accurate (so long as 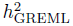 is not near either boundary) (Table 2). The fact that the GCTA standard errors are slightly larger than the corresponding empirical standard deviations of 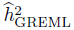 is not necessarily a drawback of GCTA because the empirical standard deviations do not reflect the variation in realized 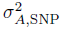 (attributable to variation in **Z**) from sample to sample. The figures of Speed et al (2012) and Zhou et al (2013) show very large standard deviations of heritability estimates in the case of few nonzeros because they varied the identities of the nonzeros across replicates. Across repeated studies of the same phenotype, where of course the identities of the nonzeros do not vary, it appears that GREML procedures produce robust standard errors after all.

**Table 2.** Comparison of empirical and GCTA standard errors

| SNP $i$ | standard deviation<br>of $\hat{h}_{\text{GREML}}^2$ | mean of GCTA<br>standard errors |
| --- | --- | --- |
| *very weakly tagged nonzeros* |  |  |
| rs4674229 | 0.0456 | 0.0637 |
| rs4716447 | 0.0539 | 0.0661 |
| rs4968679 | 0.0479 | 0.0672 |
| *weakly tagged nonzeros* |  |  |
| rs11039838 | 0.0593 | 0.0679 |
| rs4912830 | 0.0577 | 0.0681 |
| rs2654534 | 0.0577 | 0.0680 |
| *moderately tagged nonzeros* |  |  |
| rs7620645 | 0.0613 | 0.0691 |
| rs12692474 | 0.0588 | 0.0684 |
| rs4870308 | 0.0624 | 0.0691 |
| *strongly tagged nonzeros* |  |  |
| rs16841231 | 0.0596 | 0.0667 |
| rs6424728 | 0.0657 | 0.0678 |
| rs328890 | 0.0572 | 0.0660 |
| *very strongly tagged nonzeros* |  |  |
| rs10138824 | 0.0014 | 0.0596 |
| rs2718306 | 0.0073 | 0.0606 |
| rs8006587 | 0.0075 | 0.0603 |

## Discussion

In the present work, we have deduced a necessary condition for equality between the parameter 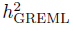 (which is estimated by software packages such as GCTA) and 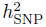 (the proportion of the phenotypic variance attributable to SNPs assayed by the given genotyping chip), in a regime allowing the inverse of the matrix **V** to be approximated by an explicit expression. In short, the condition is that 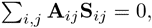 where **A***_ij_* is the realized relatedness of individuals *i* and *j* and **S***_ij_* is the deviation of the expected phenotypic product of individuals *i* and *j* from 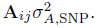 This condition in turn requires a zero correlation between relatedness and **S***_ij_*.

It is extremely plausible that this condition continues to be necessary for 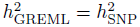 outside of the small-(*n/p*), small-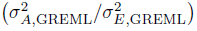 regime in which we derived it. For suppose that the condition fails, perhaps because **A***_ij_* and **S***_ij_* are positively correlated. Inspection of Eq. 7 shows that a positive correlation and consequent non-vanishing of the additional two terms causes the average phenotypic product of pairs exhibiting a given positive relatedness **A***_ij_* to exceed 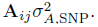 This excess phenotypic similarity between positively related individuals should “trick” GREML into overestimating heritability 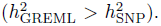 A positive (negative) correlation between relatedness and at least Term 2 in Eq. 7 can be induced by an overrepresentation of the nonzeros among the most (least) strongly tagged markers, and our simulations using small *n/p* but large 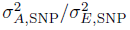 confirmed that such overrepresentation leads to a biasing of estimates in the expected direction.

This account appears to be consistent with all simulation studies of GREML performance that have appeared thus far. Speed et al (2012, 2013) also found that the extent to which nonzeros are in strong (weak) LD with neighbors is associated with the degree to which GREML produces upwardly (downwardly) biased estimates of 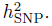 In simulations with independent markers, where of course each nonzero is as well tagged as any other marker, Zaitlen and Kraft (2012) found that GREML produces unbiased estimates of 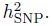 Speed et al (2012, 2013), Zhou et al (2013), Browning and Browning (2013), and Lee et al (2013) used real genetic data characterized by LD in their simulations, and we replicated their findings that choosing a large and random sample of markers to serve as the nonzeros leads to an absence of substantial bias. We note again that many of the simulations by ourselves and others finding that GREML is unbiased when 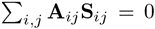 have not adhered to small *n/p* and 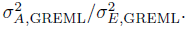 For example, Zaitlen and Kraft (2012) found that GREML can be unbiased even in the case that *n > p*. The restrictive assumptions of small *n/p* and 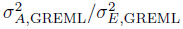 that we employed to derive the condition 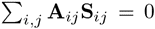 thus appear to be matters of mathematical convenience only. Therefore studies that partition heritability among different parts of the genome or that analyze highly heritable phenotypes should be sound, so long as the condition 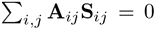 is satisfied.

We have not discussed the potential for a correlation between Terms 1 and 3 (Eq. 7). Presumably such a correlation might arise if the phenotype is subject to assortative mating, which tends to induce positive LD between causal variants (Fisher, 1918; Crow and Kimura, 1970; Lynch and Walsh, 1998). In this case individual *i*'s genotype at marker *k* is a weak proxy for *i*'s genotype at ℓ, and the fact that *i* and *j* show a positive realized similarity at *k* (Term 1) may also mean that they tend to show similarity at markers *k* and *ℓ* (Term 3). However, because phenotypes subject to assortative mating are probably also subject to natural selection, which tends to induce negative LD (Lande, 1977; Bulmer, 1980), it may be that even assortative mating does not suffice to induce a correlation between Terms 1 and 3 large enough to invalidate GREML heritability estimates. This is perhaps an important issue for further research.

Here we provide a sketch of how our account generalizes to the SNP-based genetic correlation between two traits. If we use 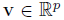 to denote the vector of partial regression coefficients in the regression of the *second* phenotype on standardized genotypes, then we can write

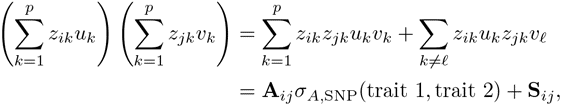

where the definition of **S***_ij_* is more or less retained from the univariate case. One complication is that the part of **S***_ij_* corresponding to LD among distinct causal loci (Term 3) is defined to be part of the genetic correlation by some authors (Lynch and Walsh, 1998). If we ignore this complication and assume LE among causal loci, then the SNP-based genetic covariance will be estimated without bias by GREML if the markers that are nonzeros with respect to both traits are an effectively random sample of all genotyped markers. Furthermore, the GREML-estimated genetic correlation (the ratio of genetic covariance to the square roots of the genetic variances) may be close to unbiased as an estimate of the true genetic correlation under fairly general conditions, since biases attributable to missing causal variants and unrepresentative tagging of nonzeros may cancel from both the numerator and denominator (Trzaskowski et al, 2013). These issues may also be a worthwhile focus of future research.

Our explication shows that GREML estimates of SNP-based heritability will be reasonably accurate under much wider circumstances than those under this approach has been previously derived. Such estimates are insensitive to the number, MAF spectrum, and coefficient magnitudes *per se* of the markers with nonzero regression coefficients. The sensitivity to the LD properties of the genomic regions containing the nonzeros, however, does raise some concern. This is the crucial question: how likely is it that the nonzeros of a given phenotype are effectively like a large and random sample of all genotyped (or imputed) markers with respect to tagging?

So far we have two sources of guidance. First, it has been empirically found that the frequency spectrum of nonzeros tends to be skewed toward low MAF (Park et al, 2011). Such a skew is also plausible for evolutionary reasons. A trait-affecting mutation is likely to face a slight selection pressure that disfavors its frequency increase (Eyre-Walker, 2010), and because a causal variant can only be in strong LD with a marker if the two sites have similar MAFs (Wray et al, 2011), the spectrum of markers tagging causal variants should also be skewed toward low MAF. Since low-MAF variants tend to be less strongly tagged on the whole, we might then expect 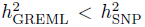 to be typical. On the other hand, because the correlation between MAF and tagging is less than perfect (∼ .30 in our larger dataset), it might be reasonable to expect that such a bias will usually be mild if the number of nonzeros is large. In the simulations of Speed et al (2012, 2013) and Lee et al (2013), even a strong correlation between MAF and coefficient magnitude introduced biases of only about .05, and perhaps underestimates of 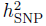 to this extent have no practical bearing on the discussion of missing heritability. Second, Speed et al (2012) have implemented an ingenious method in the LDAK package that weights markers by the extent to which they are tagged by neighbors when calculating realized relatedness. It appears from their simulations that using the resulting LD-adjusted **A** matrix to estimate 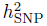 is usually successful in removing most of any bias affecting an estimate based on the unadjusted **A** matrix (Eq. 5). When they applied the LDAK method to several real phenotypes, they found a tendency for the LDAK-corrected estimates to be larger than the standard GREML estimates (supporting the notion that causal loci tend to reside at low MAF), but these increases were modest and perhaps of little practical relevance to the issue of missing heritability.

In future studies we recommend employing both the LDAK and standard GREML methods and considering their results together. Although LDAK can markedly attenuate substantial biases affecting standard GREML estimates, in some cases LDAK introduces a small bias that is otherwise absent (Speed et al, 2012, 2013; Lee et al, 2013). Perhaps surprisingly, given the mathematical nonequivalence of what we have called 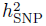 and 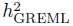, the GREML method is quite robust. It should remain a valuable tool in quantitative genetics and gene-trait mapping research for some time to come.

## Web Resources

Genome-wide Complex Trait Analysis (GCTA), http://www.complextraitgenomics.com/software/gcta

Linkage-Disequilibrium Adjusted Kinships (LDAK), http://dougspeed.com/ldak

PLINK, http://pngu.mgh.harvard.edu/∼purcell/plink, https://www.cog-genomics.org/plink2

## Acknowledgements

We thank Doug Speed and Xiang Zhou for answering our queries. This work was supported by the Intramural Program of the NIH, The National Institute of Diabetes and Digestive and Kidney Diseases (NIDDK). Assistance with phenotype harmonization and genotype cleaning, as well as with general study coordination, was provided by the Gene Environment Association Studies, GENEVA Coordinating Center (U01 HG004446). Assistance with data cleaning was provided by the National Center for Biotechnology Information.

